# *Yersinia canariae* sp. nov., isolated from a human yersiniosis case

**DOI:** 10.1101/803825

**Authors:** Scott V. Nguyen, David R. Greig, Daniel Hurley, Yu Cao, Evonne McCabe, Molly Mitchell, Claire Jenkins, Séamus Fanning

## Abstract

A Gram-negative rod from the *Yersinia* genus was isolated from a clinical case of yersiniosis in the United Kingdom. Long read sequencing data from an Oxford Nanopore Technology (ONT) MinION in conjunction with Illumina HiSeq reads were used to generate a finished quality genome of this strain. Overall Genome Related Index (OGRI) of the strain was used to determine that it was a novel species within *Yersinia*, despite biochemical similarities to *Yersinia enterocolitica*. The 16S ribosomal RNA gene accessions are MN434982-MN434987 and the accession number for the complete and closed chromosome is CP043727. The type strain is CFS3336^T^ (=<u>NCTC 14382</u>^T^/ =LMG *Accession under process*).

## INTRODUCTION AND BACKGROUND

The majority of species within the *Yersinia* genus are considered to be non-pathogenic and are found broadly within the environment. Pathogenic members of *Yersinia* have been shown to evolve independently following the acquisition of virulence genes in select lineages (1). *Yersinia pestis*, arguably one of the most historically serious zoonotic pathogens reported (2), is the agent of bubonic, pneumonic, and septicaemic plague (3). Two other species, *Yersinia enterocolitica* and *Yersinia pseudotuberculosis*, are the aetiological agents of the human gastrointestinal infection yersiniosis (4). As detailed in a 2016 report by the European Food Safety Authority, yersiniosis is the third most commonly reported zoonotic pathogen in Europe (5). *Yersinia* lends its namesake to the *Yersiniaceae* family within the *Enterobacteriales* order (6) and is comprised of 19 species, including the aforementioned pathogenic species and *Y. aldovae*, *Y. aleksiciae*, *Y. bercovieri*, *Y. entomophaga*, *Y. frederiksenii*, *Y. hibernica*, *Y. intermedia*, *Y. kristensenii*, *Y. massiliensis*, *Y. mollaretii*, *Y. nurmii*, *Y. pekkanenii*, *Y. rohdei*, *Y. ruckeri*, *Y. similis*, and *Y. wautersii*. Two additional subspecies are described with *Y. enterocolitica* subsp. *palearctica* and the recently characterised *Y. kristensenii* subsp. *rochesterensis* (7).

At a local NHS frontline hospital laboratory, diarrhoeic stool samples were tested with the GI PCR screening test (Fast-Track Diagnostics Bacterial gastroenteritis panel FTD-14.1-64 supplied by Launch Diagnostics). Specimens which tested positive for *Yersinia* by PCR were then cultured on Cefsulodin irgasan (triclosan) novobiocin (CIN) agar at 28° C for 48 hours. Isolates were referred to the Gastrointestinal Bacteria Reference Unit (GBRU) in Public Health England (PHE) for further speciation and characterisation. As per statutory reporting requirements for infectious diseases, the laboratory reported confirmed cases (PCR and/or culture positive) to PHE’s Second Generation Surveillance System (SGSS).

One such suspected *Yersinia* isolate, denoted NCTC 14382^T^, was isolated from an adult human female in the United Kingdom after travel to the Canary Islands in 2018 (8). Identification of *Yersinia* to the species level by traditional biochemical methods is difficult due to heterogeneous biochemical phenotypes (9), thus all *Yersinia* isolates receipted at the GBRU are routinely sequenced via an Illumina HiSeq 2500 and subsequent bioinformatics speciation is based on a k-mer (18-mer) based approach comparing k-mers to a known reference database (8). The closest match for this isolate by number of k-mers in the database was *Yersinia enterocolitica*. Whole genome sequencing data characterised this isolate as ST333, utilising the multi-locus sequence type (MLST) scheme developed by Hall *et al*. (8,10). A phylogenetic tree previously revealed that the isolate did not cluster with *Y. enterocolitica* instead being located on a distinct branch (8). This study aims to resolve the taxonomic placement of this isolate as a new species within *Yersinia* with the support of genomic and biochemical data. As NCTC 14382^T^ was associated with travel to the Canary Islands, the name *Yersinia canariae* sp. nov. is proposed.

## BIOCHEMICAL TESTS

API 20E strips were used in determining the phenotypes of *Y. canariae* NCTC 14382^T^ and two closely related *Yersinia* species through a gallery of biochemical tests. The latter is suitable for the identification of *Yersinia* species (11) but is dependent on incubation temperatures (12,13). Biological triplicates of the test *Yersinia* strains were assayed with API 20E test strips according to manufacturer’s instructions and incubated at both 28° and 37° C for 24 hours (**Table 1**). Incubation of NCTC 14382^T^ at 28° C revealed this strain was capable of fermenting almost all carbon sources, similar to *Y. enterocolitica* 8081 (**Fig. 2**). *Y. canariae* NCTC 14382^T^ was negative for utilisation of ODC at 28° C in contrast to *Y. enterocolitica*. Inoculated API 20E strips incubated at 37° C produced more contrasting phenotypes notably with the loss of inositol fermentation for NCTC 14382^T^ when compared to *Y. enterocolitica* (**Fig. 3**). *Y. canariae* NCTC 14382^T^ was positive for ONPG utilisation in contrast to the other two strains tested at 37° C.

**Table 1.** API 20E results for *Y. enterocolitica* 8081 (1), *Y. canariae* NCTC 14382^T^ (2), and *Y. hibernica* CFS1934 (3) at 28 °C and 37 °C. + = positive, − = negative, V = variable.

|  | 28 °C |  |  | 37 °C |  |  |
| --- | --- | --- | --- | --- | --- | --- |
|  | 1 | 2 | 3 | 1 | 2 | 3 |
| ONPG | + | + | - | V | + | - |
| ADH | - | - | - | - | - | - |
| LDC | - | - | - | - | - | - |
| ODC | + | - | - | + | - | - |
| CIT | - | - | - | - | - | - |
| H<sub>2</sub>S | - | - | - | - | - | - |
| URE | + | + | + | + | + | + |
| TDA | - | - | - | - | - | - |
| IND | + | + | - | + | + | - |
| VP | - | - | - | - | - | - |
| GEL | - | - | - | - | - | - |
| GLU | + | + | + | + | + | + |
| MAN | + | + | + | + | + | + |
| INO | + | + | + | + | - | - |
| SOR | + | + | + | + | + | + |
| RHA | - | - | - | - | - | - |
| SAC | + | + | - | + | + | - |
| MEL | - | - | - | - | - | - |
| AMY | + | + | + | + | + | - |
| ARA | + | + | + | + | + | + |

## GENOME FEATURES

*Y. canariae* NCTC 14382^T^ was previously sequenced by an Illumina HiSeq 2500 at Public Health England using the Nextera XP library preparation kit following a retrospective study on yersiniosis isolates cultured from patients between April 2004 and March 2018 (8). To generate a finished quality genome, NCTC 14382^T^ was grown in Luria-Bertani (LB) broth (Sigma) for 18 h at 25° C. Genomic DNA was extracted with a Wizard Genomic DNA Purification Kit (Promega) following the recommended manufacturer’s protocol. DNA was sequenced on the ONT MinION R9.4 flowcell (FLO-MIN106) for approximately 16 hours.

For ONT MinION data, the run metrics were inspected using NanoPlot (version 1.0) (14) before raw FAST5 files were base-called using Guppy (version 3.2.2) with the high accuracy model to FASTQ files. Adapters were trimmed from the raw reads by Porechop (version 0.2.4) using default parameters for SQK-RAD004 before the genome was *de novo* assembled with Flye (version 2.5) (15,16). The best assembly parameters were empirically determined to include the option flags “meta” and “plasmid” with coverage reduced to 30X for initial contig assembly based on a predicted genome size of ~4.73 Mbp as informed by *de novo* assembly of short read Illumina data (17). This produced a single contiguous chromosome for which the final consensus sequence was determined following four iterative rounds of long read polishing with Racon (version 1.4.3) (18) using the high accuracy base-called reads produced by Guppy that were previously adaptor trimmed by Porechop. A final round of consensus sequence correction was performed with the same long read data using Medaka (version 0.8.2). Lastly, short read Illumina data were aligned using minimap2 (version 2.17) (19) producing BAM files that were sorted and indexed with Samtools (version 1.9) (20) before four iterative rounds of short read polishing with Pilon (version 1.23) (21).

After assembly, a circular finished quality chromosome of 4,7101,54 bp devoid of any plasmid was generated. Annotation by the NCBI Prokaryotic Genome Annotation Pipeline (PGAP) identified 4,370 genes of which 4,132 were coding. 8 copies of the 5S rRNA genes, 7 copies of the 16S rRNA genes, 7 copies of the 23S rRNA genes, 81 tRNA genes, and 6 non-coding RNA genes were present, resulting in a total of 109 RNA genes.

As the genome was corrected by multiple rounds with long read aware polishing, the most common 16S rRNA gene allele (Accession: MN434982) was extracted from the genome for analysis. The full length 16S rRNA gene was queried in the EzBioCloud 16S database as previously recommended by Chun *et al*. (**Table 2**) (22,23). For much of the type strains of *Yersinia* species, NCTC 14382^T^ showed at least 98.7% or higher 16S rRNA gene similarity and thus was queried for overall genomic related index (OGRI).

**Table 2.** 16S rRNA gene pairwise similarity of *Y. canariae* (Accession: MN434982) to other type strains of *Yersinia*. Nucleotide mismatch is based on differences between the *Y. canariae* 16S rRNA gene to reference sequences.

| **Name** | **Strain** | **Accession** | **Pairwise Similarity (%)** | **Mismatch /Total nt** |
| --- | --- | --- | --- | --- |
| *Yersinia frederiksenii* | ATCC 33641 | JPPS01000006 | 99.11 | 13/1465 |
| *Yersinia pekkanenii* | CIP 110230 | CWJL01000114 | 98.91 | 16/1465 |
| *Yersinia hibernica* | CFS1934 | MK129259 | 98.91 | 16/1465 |
| *Yersinia kristensenii* subsp. *rochesterensis* | EPLC-04 | KJ606916 | 98.90 | 16/1457 |
| *Yersinia aldovae* | ATCC 35236 | AF366376 | 98.84 | 17/1461 |
| *Yersinia rohdei* | ATCC 43380 | ACCD01000072 | 98.77 | 18/1465 |
| *Yersinia enterocolitica* subsp. *polarctica* | Y11 | FR729477 | 98.77 | 18/1465 |
| *Yersinia massiliensis* | CCUG 53443 | CAKR01000050 | 98.77 | 18/1465 |
| *Yersinia nurmii* | CIP 110231 | CPYD01000031 | 98.77 | 18/1465 |
| *Yersinia intermedia* | ATCC 29909 | AF366380 | 98.77 | 18/1461 |
| *Yersinia entomophaga* | MH96 | CP010029 | 98.70 | 19/1464 |
| *Yersinia mollaretii* | ATCC 43969 | AF366382 | 98.70 | 19/1461 |
| *Yersinia kristensenii* subsp. *kristensenii* | ATCC 33638 | ACCA01000078 | 98.63 | 20/1465 |
| *Yersinia bercovieri* | ATCC 43970 | AF366377 | 98.56 | 21/1461 |
| *Yersinia pestis* | NCTC 5923 | AF366383 | 98.43 | 23/1461 |
| *Yersinia pseudotuberculosis* | NBRC 105692 | BAUR01000130 | 98.36 | 24/1465 |
| *Yersinia similis* | Y228 | CP007230 | 98.29 | 25/1465 |
| *Yersinia wautersii* | 12-219N1 | HG326166 | 98.29 | 25/1464 |
| *Yersinia aleksiciae* | DSM 14987 | CP011975 | 98.16 | 27/1465 |
| *Yersinia enterocolitica* subsp. *enterocolitica* | ATCC 9610 | JPDV01000006 | 98.02 | 29/1465 |
| *Yersinia ruckeri* | ATCC 29473 | JPPT01000003 | 97.82 | 32/1465 |

The average nucleotide identity (ANI) of NCTC 14382^T^ was determined by FastANI (v1.2) against the type sequences of all other *Yersinia* species (24). *Y. canariae* NCTC 14382^T^ was most closely related to *Y. hibernica*, *Y. enterocolitica* subsp. *enterocolitica*, *Y. enterocolitica* subsp. *palearctica*, *Y. kristensenii* subsp. *rochesterensis*, and *Y. kristensenii* subsp. *kristensenii* based on ANI values (**Table 3**). However, the ANI values for NCTC 14382^T^ are below the threshold of ≤95% ANI when compared to the type strains of other *Yersinia* species, thus suggesting taxonomic placement of NCTC 14382^T^ into a novel species. The digital DNA-DNA hybridization (dDDH) values were calculated with the Type (Strain) Genome Server hosted by the Deutsche Sammlung von Mikroorganismen und Zellkulturen (DSMZ) (25). Using the recommended formula d_4_ (25,26) with the BLAST+ local alignment tool, *in silico* DNA-DNA hybridization values of NCTC 14382^T^ revealed additional OGRI data for the support of a novel species (**Table 3**).

**Table 3.**
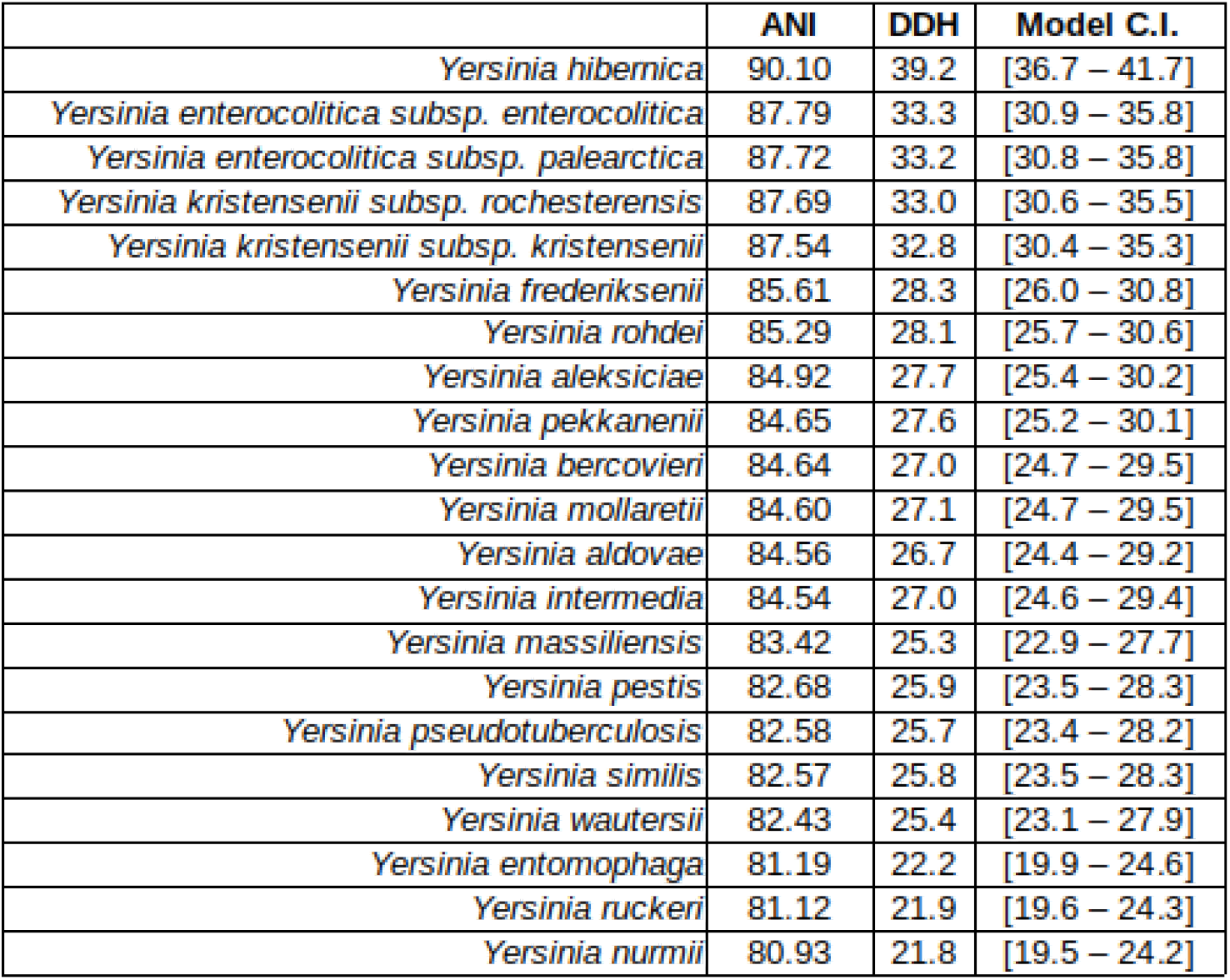
ANI and DDH (d_4_ formula) of *Y. canariae* NCTC 14382^T^ queried against type strains of other *Yersinia* species. Closely related *Y. hibernica* was at 90.10% ANI and 39.2% DDH, well under the 95% ANI and 70% DDH cutoff proposed by Chun *et al.* (22).

A phylogenetic tree based on conserved core sequences used in a whole genome alignment was generated by Parsnp as previously described (27). The resulting phylogenetic tree was visualized in EvolView2 (28) and showed that *Y. canariae* clustered distinctly from other *Yersinia* species and was most closely related to the newly described *Y. hibernica* (27) (**Fig. 1**).

**Figure 1.**
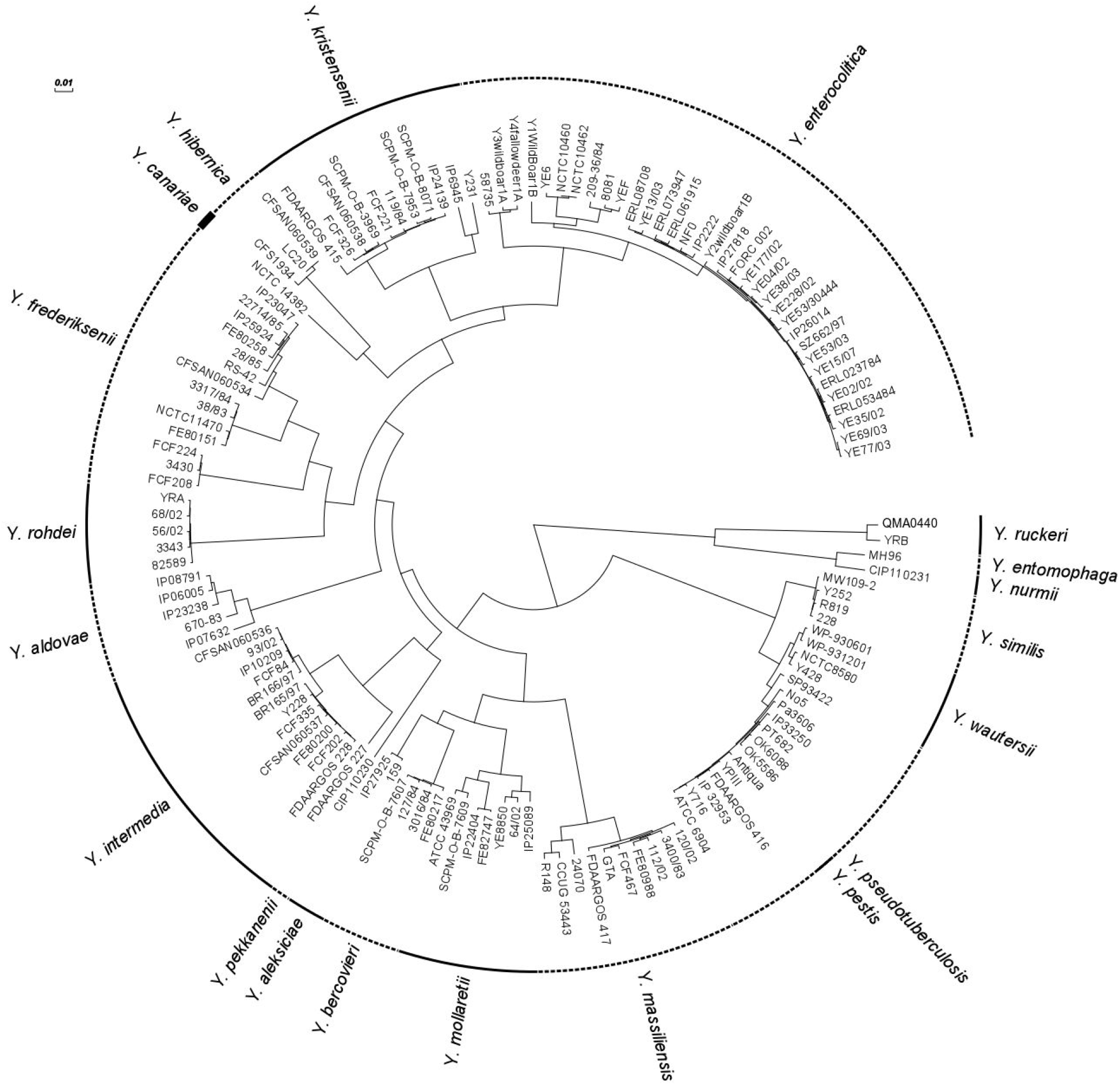
Taxonomic placement of *Y. canariae* NCTC 14382^T^ in relation to other *Yersinia* species on the basis of whole genome alignment of conserved core sequences.

**Figure 2.**
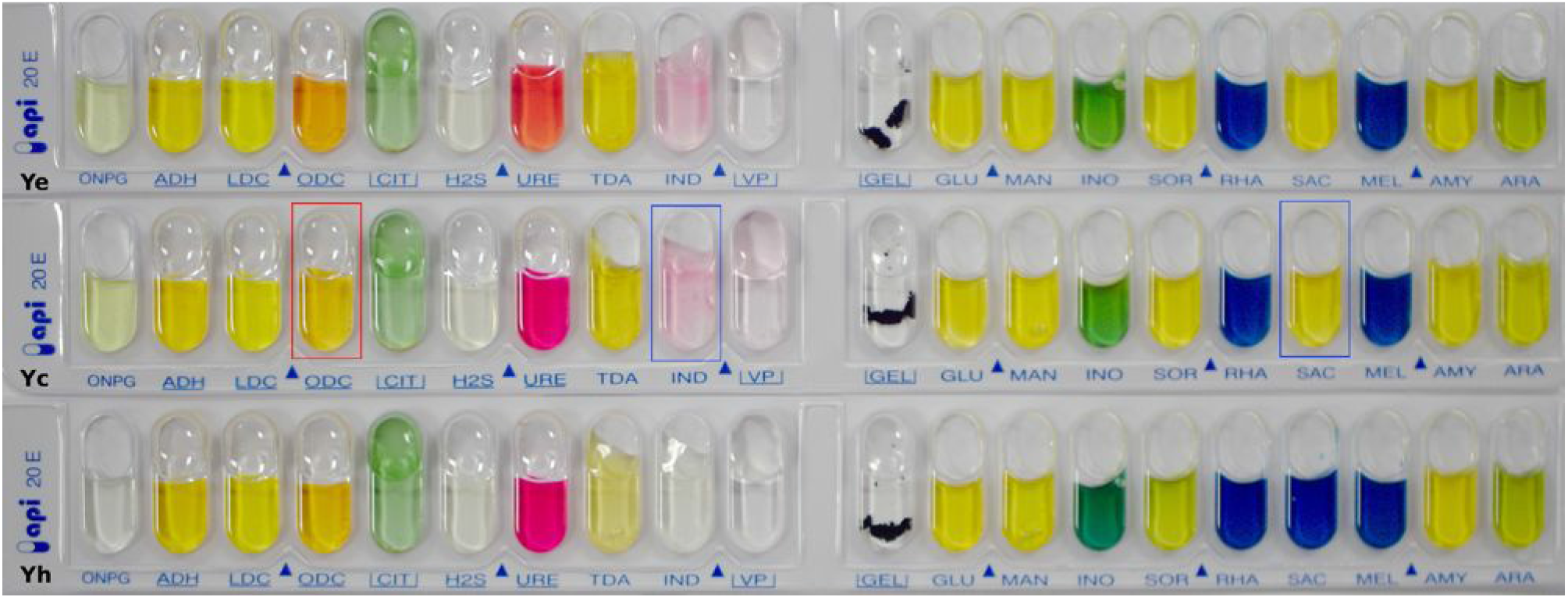
Growth of *Y. enterocolitica* 8081 (**Ye**), *Y. canariae* NCTC 14382^T^ (**Yc**), and *Y. hibernica* CFS1934 (**Yh**) at 28 °C after 20 hours. Boxes highlight biochemical features distinct from *Y. enterocolitica* (red) or *Y. hibernica* (blue).

**Figure 3.**
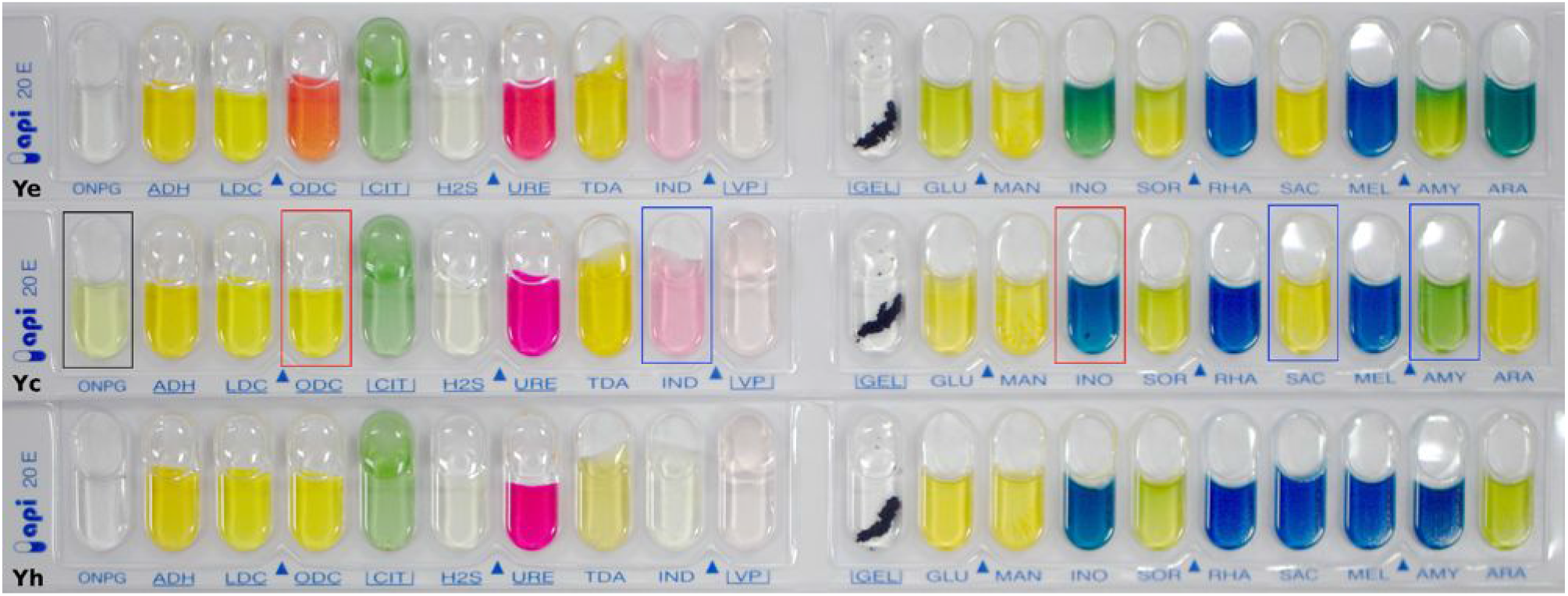
Growth of *Y. enterocolitica* 8081 (**Ye**), *Y. canariae* NCTC 14382^T^ (**Yc**), and *Y. hibernica* CFS1934 (**Yh**) at 37 °C after 20 hours. Boxes highlight biochemical features distinct from *Y. enterocolitica* (red) or *Y. hibernica* (blue) or both (black).

On the basis of the biochemical profile, phylogenetic relationships, and OGRI data of isolate NCTC 14382^T^, evidence for a new species within *Yersinia* is conclusive in which the name *Yersinia canariae* sp. nov. is proposed.

### Description of *Yersinia canariae* sp. nov

*Yersinia canariae* (ca.na’ri.ae. N.L. gen. n. *canariae* from the Canary Islands in which this strain was associated with travel to the Canary Islands, *Canariae insulae*).

Cells grow aerobically at 25-37 °C on LB agar, producing 1.5-2.0 mm diameter colonies after 24 hours. At 28 °C, cells are positive for ortho-nitrophenyl-β-D-galactopyranoside hydrolysis, urease utilisation, indole production and fermentation of D-glucose, D-mannitol, inositol, D-sorbitol, D-sucrose, amygdalin, and L-arabinose. At 28 °C, cells are negative for utilisation of L-arginine, L-lysine, L-ornithine, trisodium citrate, H_2_S production, tryptophan deaminase, gelatinase, L-rhamnose fermentation, and D-melibiose fermentation. In contrast to growth at 28 °C, cells do not ferment inositol at 37 °C. The DNA G+C content is 47.2% and the chromosomal length of the type strain is 4710154 bp.

The type strain, CFS3336^T^ (=NCTC 14382^T^ =LMG *Accession under process*), was isolated in the United Kingdom from a yersiniosis case associated with travel to the Canary Islands. The complete genome of NCTC 14382^T^ has been deposited into GenBank (accession number, CP043727).

## REFERENCES

1. Reuter S, Connor TR, Barquist L, Walker D, Feltwell T, Harris SR, et al. Parallel independent evolution of pathogenicity within the genus *Yersinia*. Proc Natl Acad Sci USA. 2014 May 6;111(18):6768–73.

2. Bramanti B, Dean KR, Walløe L, Chr Stenseth N. The Third Plague Pandemic in Europe. Proc Biol Sci. 2019 Apr 24;286(1901):20182429.

3. Atkinson S, Williams P. *Yersinia* virulence factors - a sophisticated arsenal for combating host defences. F1000Research. 2016 Jun 14;5:F1000 Faculty Rev-1370.

4. Davis KM. All *Yersinia* Are Not Created Equal: Phenotypic Adaptation to Distinct Niches Within Mammalian Tissues. Frontiers in Cellular and Infection Microbiology. 2018;8:261.

5. European Food Safety Authority, European Centre for Disease Prevention and Control. The European Union summary report on trends and sources of zoonoses, zoonotic agents and food-borne outbreaks in 2013. EFSA Journal. 2015 Jan 1;13(1):3991.

6. Adeolu M, Alnajar S, Naushad S, S Gupta R. Genome-based phylogeny and taxonomy of the “*Enterobacteriales*”: proposal for *Enterobacterales* ord. nov. divided into the families *Enterobacteriaceae*, *Erwiniaceae* fam. nov., *Pectobacteriaceae* fam. nov., *Yersiniaceae* fam. nov., *Hafniaceae* fam. nov., *Morganellaceae* fam. nov., and *Budviciaceae* fam. nov. Int J Syst Evol Microbiol. 2016 Dec;66(12):5575–99.

7. Cunningham SA, Jeraldo P, Patel R. *Yersinia kristensenii* subsp. *rochesterensis* subsp. nov., isolated from human feces. Int J Syst Evol Microbiol. 2019 Aug;69(8):2292–8.

8. Hunter E, Greig DR, Schaefer U, Wright MJ, Dallman TJ, McNally A, et al. Identification and typing of *Yersinia enterocolitica* and *Yersinia pseudotuberculosis* isolated from human clinical specimens in England between 2004 and 2018. J Med Microbiol. 2019 Apr;68(4):538–48.

9. Fredriksson-Ahomaa M, Joutsen S, Laukkanen-Ninios R. Identification of *Yersinia* at the Species and Subspecies Levels Is Challenging. Curr Clin Micro Rpt. 2018 Jun 1;5(2):135–42.

10. Hall M, Chattaway MA, Reuter S, Savin C, Strauch E, Carniel E, et al. Use of whole-genus genome sequence data to develop a multilocus sequence typing tool that accurately identifies *Yersinia* isolates to the species and subspecies levels. J Clin Microbiol. 2015 Jan;53(1):35–42.

11. Arnold T, Neubauer H, Nikolaou K, Roesler U, Hensel A. Identification of *Yersinia enterocolitica* in minced meat: a comparative analysis of API 20E, *Yersinia* identification kit and a 16S rRNA-based PCR method. J Vet Med B Infect Dis Vet Public Health. 2004 Feb;51(1):23–7.

12. Archer JR, Schell RF, Pennell DR, Wick PD. Identification of *Yersinia* spp. with the API 20E system. J Clin Microbiol. 1987 Dec;25(12):2398–9.

13. Sharma NK, Doyle PW, Gerbasi SA, Jessop JH. Identification of *Yersinia* species by the API 20E. J Clin Microbiol. 1990 Jun;28(6):1443–4.

14. De Coster W, D’Hert S, Schultz DT, Cruts M, Van Broeckhoven C. NanoPack: visualizing and processing long-read sequencing data. Bioinformatics. 2018 Aug 1;34(15):2666–9.

15. Kolmogorov M, Yuan J, Lin Y, Pevzner PA. Assembly of long, error-prone reads using repeat graphs. Nat Biotechnol. 2019 May;37(5):540–6.

16. Lin Y, Yuan J, Kolmogorov M, Shen MW, Chaisson M, Pevzner PA. Assembly of long error-prone reads using de Bruijn graphs. PNAS. 2016 Dec 27;113(52):E8396–405.

17. Bankevich A, Nurk S, Antipov D, Gurevich AA, Dvorkin M, Kulikov AS, et al. SPAdes: A New Genome Assembly Algorithm and Its Applications to Single-Cell Sequencing. Journal of Computational Biology. 2012 May;19(5):455–77.

18. Vaser R, Sovic I, Nagarajan N, Sikic M. Fast and accurate de novo genome assembly from long uncorrected reads. Genome Res. 2017 Jan 18;gr.214270.116.

19. Li H. Minimap2: pairwise alignment for nucleotide sequences. Bioinformatics. 2018 Sep 15;34(18):3094–100.

20. Li H, Handsaker B, Wysoker A, Fennell T, Ruan J, Homer N, et al. The Sequence Alignment/Map format and SAMtools. Bioinformatics. 2009 Aug 15;25(16):2078–9.

21. Walker BJ, Abeel T, Shea T, Priest M, Abouelliel A, Sakthikumar S, et al. Pilon: An Integrated Tool for Comprehensive Microbial Variant Detection and Genome Assembly Improvement. PLOS ONE. 2014 Nov 19;9(11):e112963.

22. Chun J, Oren A, Ventosa A, Christensen H, Arahal DR, da Costa MS, et al. Proposed minimal standards for the use of genome data for the taxonomy of prokaryotes. International Journal of Systematic and Evolutionary Microbiology. 2018;68(1):461–6.

23. Yoon S-H, Ha S-M, Kwon S, Lim J, Kim Y, Seo H, et al. Introducing EzBioCloud: a taxonomically united database of 16S rRNA gene sequences and whole-genome assemblies. Int J Syst Evol Microbiol. 2017 May;67(5):1613–7.

24. Jain C, Rodriguez-R LM, Phillippy AM, Konstantinidis KT, Aluru S. High throughput ANI analysis of 90K prokaryotic genomes reveals clear species boundaries. Nat Commun. 2018 30;9(1):5114.

25. Meier-Kolthoff JP, Göker M. TYGS is an automated high-throughput platform for state-of-the-art genome-based taxonomy. Nat Commun. 2019 16;10(1):2182.

26. Meier-Kolthoff JP, Auch AF, Klenk H-P, Göker M. Genome sequence-based species delimitation with confidence intervals and improved distance functions. BMC Bioinformatics. 2013;14:60–60.

27. Nguyen SV, Muthappa DM, Hurley D, Donoghue O, McCabe E, Anes J, et al. *Yersinia hibernica* sp. nov., isolated from pig-production environments. Int J Syst Evol Microbiol. 2019 Jul;69(7):2023–7.

28. He Z, Zhang H, Gao S, Lercher MJ, Chen WH, Hu S. Evolview v2: an online visualization and management tool for customized and annotated phylogenetic trees. Nucleic Acids Res [Internet]. 2016;44(Web Server Issue): W236–W241.

